# Network-based protein-protein interaction prediction method maps perturbations of cancer interactome

**DOI:** 10.1101/2020.07.01.181776

**Authors:** Jiajun Qiu, Kui Chen, Chunlong Zhong, Sihao Zhu, Xiao Ma

## Abstract

The perturbations of protein-protein interactions (PPIs) were found to be the main cause of cancer. Previous gene relationship prediction methods which were trained with non-disease gene interaction data were not compatible to map the PPI network in cancer. Therefore, we established a novel cancer specific PPI prediction method dubbed NECARE, which was based on relational graph convolutional network (R-GCN) with knowledge-based features. It achieved the best performance with a Matthews correlation coefficient (MCC) = 0.84±0.03 and an F1 = 91±2% comparing with other methods. With NECARE, we mapped the cancer interactome atlas and revealed that the perturbations of PPIs were enriched on 1362 genes, which were named cancer hub genes. Those genes were found to over-represent with mutations occurring at protein-macromolecules binding interfaces. Furthermore, over 42% of cancer treatment-related genes belonged to hub genes, which were significantly related to the prognosis of 32 types of cancers. Finally, by coimmunoprecipitation, we confirmed that the NECARE prediction method was highly reliable with a 90% accuracy. Overall, we provided the novel network-based protein-protein interaction prediction method and mapped the perturbation of cancer interactome. NECARE is available at: https://github.com/JiajunQiu/NECARE.

## Introduction

Cells are biological systems that employ a large number of genes and signaling pathways to coordinate multiple functions (1). Therefore, instead of acting in isolation, genes interact with each other and work as part of complex networks (2). The completeness of these networks is the foundation of the normal biological systems, while perturbation of them can result in the pathological state. Recent studies have already found network perturbation is the cause of cancers, rather than the dysregulation of single genes (2). Gene network in cancer can be perturbed by many factors, one of which could be mutations. Disease-causing mutations can not only produce a mutated gene and thus a mutated protein, but also disturb the interactions between the mutated protein and its normal molecular partners (3). Additionally, distinct mutations will cause different molecular defects in proteins, and they may lead to distinct perturbations of gene networks, giving rise to distinct phenotypic outcomes (4). Nonsense mutations that grossly destabilize a protein structure can be modeled as removing a protein node from the network (Fig. 1A). Alternatively, missense mutations may give rise to partially functional gene products with specific changes in distinct biophysical or biochemical interactions (Fig. 1B) (4). Furthermore, studies have already found that missense mutations in cancer are more likely to occur on the interaction interface of proteins. Thus, network perturbation, instead of single gene dysregulation, has been found to be the reason for human diseases, especially cancers (5). For example, in cancer, TP53, a well-known tumor suppressor gene (Fig. 1C), loses many interactions with other important genes, such as PTEN and MDM2 (6). However, new genes, such as CDK4, have been discovered to interact with TP53. In the normal network, the cross-talk line from TP53 to CDKN2A is TP53-MDM2-CDKN2A, but in cancer, the cross-talk line is TP53-CDK4-STK11-CDKN2A (7). Therefore, in cancer, mutations lead to reconstruction of the gene network rather than the simple destruction, making the gene network in cancer tissues very different from that in normal tissues.

**Fig. 1:**
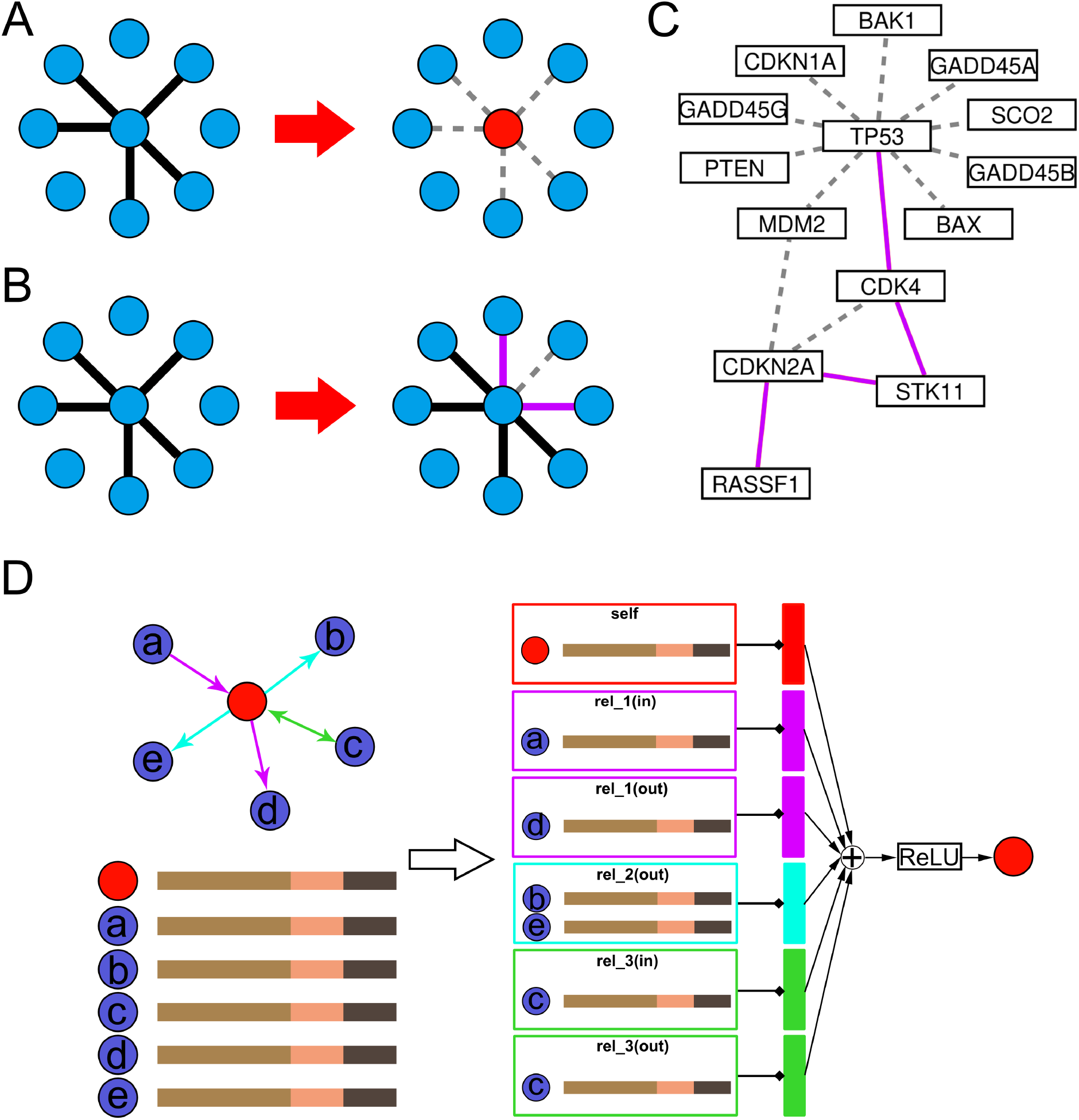
Illustration of the perturbation of the gene relationship network and NECARE algorithm. **<u>Panel A-C</u>** introduce the concept of gene network perturbation. (A) Each node represents a gene. Mutations such as nonsense mutations could cause the node to be totally inactive or absent (red) and lose all the edges connected to this node (gray dashed edges). (B) Each node represents a gene. Mutations such as missense mutations could cause the gain or loss of specific edges (purple edges mean the new gained edges due to the mutations; gray dashed edge means lost interaction), while the center node is not totally inactive. (C) This is an example of the perturbation of the gene relationship network in cancer. The example is based on the KEGG database (6). Gray dashed edges are the interactions that are lost in cancer, and purple edges are the new interactions in which genes are involved in cancer. **<u>Panel D</u>** is a simple example to show how we represent the gene (red node) by NECARE with R-GCN. Nodes a-e and the red node represent different genes, and the red node is set as the target gene. Nodes a-e are all in contact with the red node, and different colored edges represent different types of interactions. First, each node is represented by a feature vector that contains three parts: (tan: OPA2Vec; salmon: TCGA-based expression feature; and taupe: TCGA-based mutation feature). Then, to represent the red node, the feature vectors are gathered and transformed for each relation type individually (for both in- and out-edges; also, a self-loop is included). The resulted representation (vertical rectangles with different colours for different relationship types) is summed up and passed to an activation function (ReLU).

There have been some studies about cancer network perturbations (8). For example, James West et al. tried to identify genes with network perturbations by calculating the network entropy (9). Maxim Grechkin et al. also identified perturbed genes through inferred gene regulators and their expression (2). As these studies were based on only the coexpression of genes, their network was more likely to reflect the relationships (activation and repression) between transcriptional factors and their targets. However, these studies failed to consider physical relationships such as protein-protein interactions (PPIs), which are also important types of gene relationships and are significantly different from coexpression networks based on topological comparisons (10).

To date, there already exist different kinds of PPIs prediction methods for non-disease situation and they fall into three categories: 1) **Structure-based methods**, which are based on the 3D structure of proteins and limited to proteins with PDB structures (11). Structure-based methods are better at predicting physical interactions. 2) **Sequence-based prediction methods,** which attempt to predict interactions by the sequences of two candidate proteins (12). 3) **Network-based methods** that predict interactions based on the known network. Unlike other methods which only consider two candidate proteins, network-based methods also consider their known neighbors (13).

In our study, we established a novel cancer PPI prediction method, dubbed NECARE (**ne**twork-based **ca**ncer gene **re**lationship prediction), to investigate the whole cancer gene PPI map. Here we applied a relational graph convolutional network (R-GCN) with knowledge-based features. One crucial novelty of this work is that, unlike previous network-based node relationship prediction algorithms, NECARE considers the type and direction of gene links at the input space, and it can also distinguish the direction of the output (predicted gene interaction). And, NECARE was found to outperform the other algorithms (both network- and sequence-based algorithms) in predicting cancer PPIs. Thus, our tool can help other researchers to determine the possible upstream and downstream molecular partners of their target genes in cancer.

Furthermore, we mapped the cancer interactome and analyzed the perturbations of PPIs in cancer with NECARE. We found that the PPI perturbations were enriched in some specific genes that were defined as cancer hub genes in our study. These hub genes were significantly related to the prognosis of 32 types of cancers. Many of these hub genes have already been well studied in previous cancer studies or served as drug targets. These results indicated that our results can potentially provide the targets of future cancer studies. Finally, we selected 20 pairs of PPI and verified the interaction of 18 pairs by coimmunoprecipitation, which demonstrated that NECARE prediction method was highly reliable with a 90% accuracy.

## Methods

### General gene network data

To build a knowledge graph which contained knowledge of the relationship between genes for R-GCN, we extracted the general gene network data from the following three databases: 1) STRING (14), a famous database for known protein-protein associations, from which we extracted data about the experimental annotated human protein-protein associations; 2) Kyoto Encyclopedia of Genes and Genomes (KEGG) (6), a well-known publicly accessible pathway database, from which we extracted human non-disease pathway; and 3) HIPPIE (15), which contains experimentally detected PPIs from IntAct (16), MINT (17), BioGRID (18), HPRD (19), DIP (20), BIND (21) and MIPS (22). Overall, our general gene network data contained 551850 pairs of inetractions (Table S1). The whole dataset is available from (github.com/JiajunQiu/NECARE/dataset/NECARE.graph).

### Cancer protein-protein interaction data

Cancer protein-protein interaction data served as the training data for the R-GCN. We obtained cancer PPI data from the KEGG and Reactome databases (23), which served as the positive training set. We also included the OncoPPI database (7), which is a cancer-specific PPI database, in our positive training set. The negative training data were the pairs of relationships with “disassociation/missing interaction” or other negative annotations in the KEGG cancer related pathways.

Overall, we have 933 positive interactions (links) and 1308 negative interactions (links). The whole dataset is available from (github.com/JiajunQiu/NECARE/dataset/NECARE_TrainingData.txt).

### The 5-fold cross-validation

We applied a 5-fold cross-validation approach for the training mode (Fig. S1). Technically, we divided the training set into five parts. In each rotation, we used three of the five parts for training, one for cross-training (optimize hyperparameters, including number of hidden units in neural network, early stop, etc.), and one for testing. Overall, we optimized the parameters in the cross-training set and tested the final performance in the testing set. The testing set was never used in the hyperparameter optimization and feature selection.

### Relational graph convolutional networks

Graph convolutional networks (GCNs) can be understood as special cases of a simply differentiable message-passing framework. Information can be obtained from the neighbors of each node in the GCN. The R-GCN is an extension of the GCN (24). It accounts for the edge type and direction and can calculate the forward-pass update of an entity or node denoted in relational (directed and labeled) multigraphs (24) (Fig. 1D).

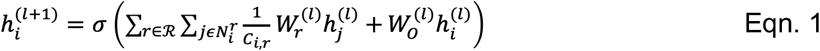

In Eqn. 1, if we define the directed and labeled multigraphs as 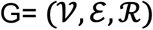 with the nodes defined as 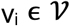, labeled edges as 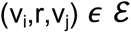, and edge type as 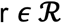, then 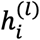 is the hidden state of node v_i_ in the i-th layer of the neural network. 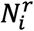 denotes the set of neighbor indices of node v_i_ under the relation 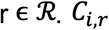 is a normalization constant, which is defined as the degree of the target node of an edge. 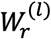 is a form of weight sharing among different relation types, and 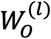 is a weight matrix for the linear message transformation. The incoming messages from neighbors are accumulated and then passed through an activation function σ such as ReLU (24). Therefore, in our study, instead of only considering the gene itself, information about each gene was obtained from other genes that contacted it.

Regarding to the feature we used to train the model, it was a combination of two parts. Part one was the OPA2Vec vector of each gene, which was a knowledge-based feature (25). OPA2Vec is a tool that can be used to produce feature vectors for biological entities from ontology. OPA2Vec used mainly metadata from the ontology in the form of annotation properties as the main source of data. In this study, we used the OPA2Vec pretrained model based on PubMed data, and the annotation file was downloaded from http://purl.obolibrary.org/obo/go.owl. Part two was the cancer-specific feature based on The Cancer Genome Atlas (TCGA), including the expression profile of each gene in 32 different types of cancer and the mutation rate among patients for each type of cancer.

### Performance evaluation

We evaluated the performance of the prediction via a variety of measures. For simplicity, we used the following standard annotations: true positives (TP) were the correctly predicted gene relationships in cancer, while false positives (FP) were the gene pairs that had no links in cancer and were incorrectly predicted to have interactions. True negatives (TN) were the correctly predicted noninteractions, and false negatives (FN) were the gene pairs that had interactions but were not a correctly predicted.

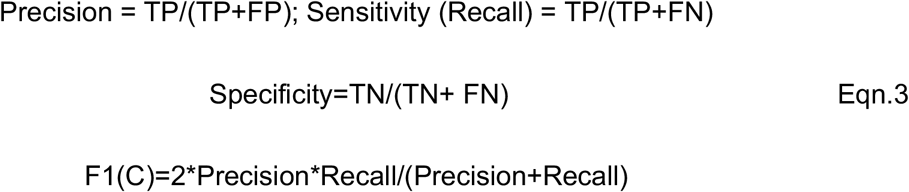

We also calculated the Matthews correlation coefficient (MCC) and area under the curve (AUC):

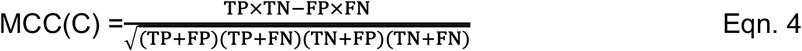

### Error estimates

Error rates for the evaluation measures were estimated by bootstrapping (without replacement to render more conservative estimates), i.e., by resampling the set of samples used for the evaluation 1000 times and calculating the standard deviation of those 1000 different results. Each of these sample sets contained 50% of the original samples (picked randomly again, without replacement).

### Cancer hub gene identification

Cancer hub genes were defined as those genes that significantly lost (or gained) links in the cancer network. We used the cancer gene links connecting with an equal likelihood at the genes in the network as a null model. We assumed that, for a particular gene (node) to be called a putative hub gene, more links (gained or lost) must connect to that gene than expected by chance if the links were randomly connected to the genes in the network. Randomly, the frequency of links connected to any particular residue followed a binomial distribution:

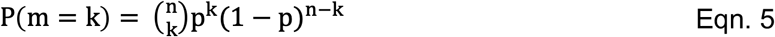

where n is the 2x total number of links, k is the number of links connecting to a particular node, p is the probability of any individual link connecting at a particular node, and P (m=k) is precisely the probability of observed k links at a single node. Since our null model assumes an equal likelihood of links at any node, we used p = 2/L, where L is the overall number of unique nodes in the network.

Thus, to assign a probability to the observation of k links connecting at a particular node by chance (i.e., a P-value), we calculated the probability of at least k links connecting at a particular node from our null model:

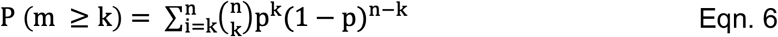

To correct for and test multiple hypotheses, the p-values for all considered hub genes were adjusted using the Bonferroni correction method.

Eigenvector centrality was a measure of the influence of a node in a network. The regular eigenvector centrality of each gene in the network was the eigenvector of the adjacency matrix with the largest unique eigenvalue. Here, in our study, we applied a variant of eigenvector centrality (26). The final centrality values followed the SoftMax probability: any node that you randomly picked up would reach a certain node in the network.

### Clinically related cancer genes

Cancer genes related to clinical treatment were downloaded from the Tumor Alterations Relevant for GEnomics-driven Therapy (TARGET) database (https://software.broadinstitute.org/cancer/cga/target). TARGET (tumor alterations relevant for genomics-driven therapy) is a database of genes that, when somatically altered in cancer, are directly linked to a clinical action. TARGET genes is associated with response or resistance to a therapy, diagnosis, and/or prognosis.

### Survival analysis of hub genes

To assess the association of hub genes with survival outcomes, we obtained the mutation and clinical prognosis data of 32 different types of cancers from the TCGA (Table S2). For each cancer, we first calculated hazard ratios (HRs) and P-values (log-rank test) for each involved gene by Cox proportional hazards regression analysis using the coxph function of the R survival package (v. 2.37.2). Then, for each cancer, we integrated the hub genes with a significant P-value (cutoff: 0.05) into a combined mutation score (MS):

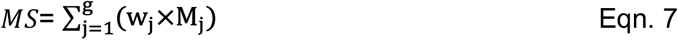

where *M_j_* is whether gene *j* is mutated in the tumor sample of the patient (1 for mutated and 0 for nonmutated) and *w_j_* is set to 1 or −1, depending on the HR of each gene (1 for HR≥1 and −1 for HR<1). The median value (50%) or the automatically selected best cutoff value of the MS was used to divide the corresponding patients into high- and low-MS groups for Kaplan–Meier analysis of their association with the 10-year survival.

## Results

### Establish network-based cancer protein-protein network prediction method (NECARE)

The PPI network in cancer is different from that in normal (non-cancer) situations. To reveal PPI network perturbation in cancer, we designed the novel network-based cancer-specific PPI prediction method: NECARE. Basically, instead of looking at only the particular gene, NECARE also obtained the information of its neighboring genes. NECARE is different from previous network-based algorithms: it accounts for the type and direction of edges at the input space (Fig. 1D).

In our study, we tested two kinds of features for the neural network: 1) ontology-based features (OPA2Vec) and 2) TCGA-based expression and mutation profiles. Their performance was compared in the cross-training set (Fig. S1). The combination of OPA2Vec and TCGA worked better than each of them alone, reaching an MCC = 0.85 (Fig. S2). Thus, the combination of OPA2Vec- and TCGA-based (expression and mutation) profiles was selected as the features for NECARE.

We also performed hyperparameter optimization in the cross-training set. Table S3 shows the hyperparameters we used in the final model of NECARE.

Finally, we evaluated the performance of NECARE in the testing set. Overall, NECARE achieved an F1 = 91±2% and an MCC = 0.84±0.03(Table S4). In addition, we also determined the reliability index (RI) of NECARE (Fig. 2A). RI was correlated with its performance and can be used to measure its prediction performance. The RI ranged from −100 to 100 (-100 meant most reliable negative prediction and 100 meant most reliable positive prediction). For instance, the subset of predictions at RI ≥0 had a precision of >90% (Fig. 2A: red line at x=0). This level covered approximately 92% of all predictions (Fig. 2A: blue line at x=0). When increasing the RI to 80 (dashed line), the precision reached 95% (Fig. 2A: red line at x=80), but it can cover only 74% of all predictions (Fig. 2A: blue line at x=80). Therefore, basically, a higher RI represented a more reliable prediction. The RI was also calculated for the negative prediction (noninteracting prediction) (Fig. 2B). At RI = 0, the precision for the negative prediction was 94%, and it increased to 97% at RI=-80 (Fig. 2B).

**Fig. 2:**
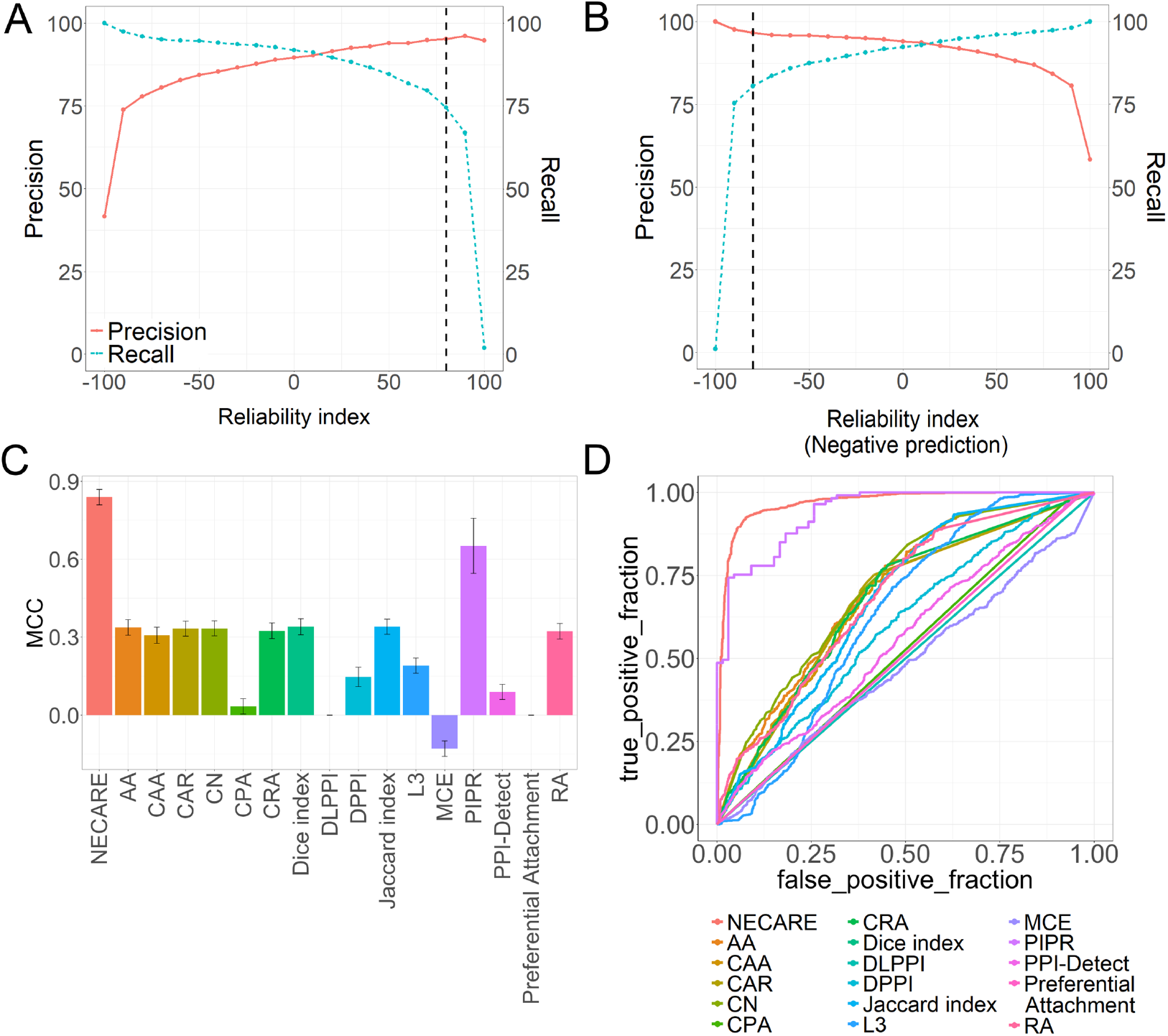
Network-based cancer gene relationship (NECARE) prediction. (A) All machine learning solutions reflect the strength of a prediction even for binary classifications. This graph relates the prediction strength to the performance. The x-axes give the prediction strength as the RI (from −100: very reliable noninteraction to 100: very reliable interaction). The y-axes reflect the precision percentage (red line, Eqn. 3) and recall percentage (blue line, Eqn. 3). The precision is proportional to the prediction strengths, i.e., predictions with a higher RI are, on average, better than predictions with a lower RI. For example, for all the gene relationship predictions with RI>80 (black dashed line), approximately 96% are correct predictions. (B) This graph relates prediction strength to performance for negative predictions (noninteractions). For example, for all the negative gene relationship predictions with RI<-80 (black dashed line), approximately 92% are correct predictions. (C) The MCC (Eqn. 4) was determined for a comparison among different methods on the test set, and our method NECARE obtains the highest MCC: 0.84. (D) ROC curve comparison for different methods based on the test set. NECARE has the largest AUC: 0.97.

### NECARE outperformed other algorithms

As NECARE is a network-based method, we first compared it with other network-based node relationship prediction algorithms such as the state of art method L3 (27), and the methods they compared in their research. We also compared NECARE with other state-of-the-art sequence-based deep learning PPI prediction methods such as PIPR (28) and DPPI (29) (Fig. 2C,2D).

First, we drew the ROC (receiver operating characteristic) curves for all the methods (Fig. 2D) and calculated the AUC for them. Our method achieved the best performance with an AUC = 0.97 (Fig. 2D, Table S4), while most of the other methods had an AUC of approximately 0.60 (Table S4).

For the detailed metrics, NECARE reached the highest F1 (91±2%) and MCC (0.84±0.03) in the comparison (Fig. 2C, Table S4). The RCNN (recurrent convolutional neural network)-based method PIPR achieved the highest precision of 94±1% (precision for NECARE was 90±2%). However, PIPR had a low specificity of 83±8% and MCC of 0.65±0.10, and the specificity of NECARE was 92±2% (Table S4).

Overall, we can conclude that NECARE is currently the best prediction method that can be used to identify gene relationships in cancer.

### Cancer hub genes discovered by NECARE

By applying NECARE, unlike previous studies that were limited to the coexpression between genes (2), we were able to reveal the comprehensive and rigorous perturbation of the cancer gene network. We mapped the cancer gene interactome with its highly reliable predictions (|RI| ≥ 0.8, Figs. 2A and 2B). On average, each gene lost 20 edges with the other genes in the cancer gene network. However, they obtained approximately 125 new edges on average (Fig. S3, red dashed lines). This indicated our hypothesis that instead of simply being fractured, the gene network in cancer is reprogrammed.

Furthermore, we assumed that the perturbation was not evenly distributed among all the genes. Some genes may hold more perturbations than others. Genes enriched with network perturbations (gained/lost links) were defined as cancer hub genes. Finally, we identified 1362 genes enriched with network perturbations in cancer (Fig. 3A, Table S5).

**Fig. 3:**
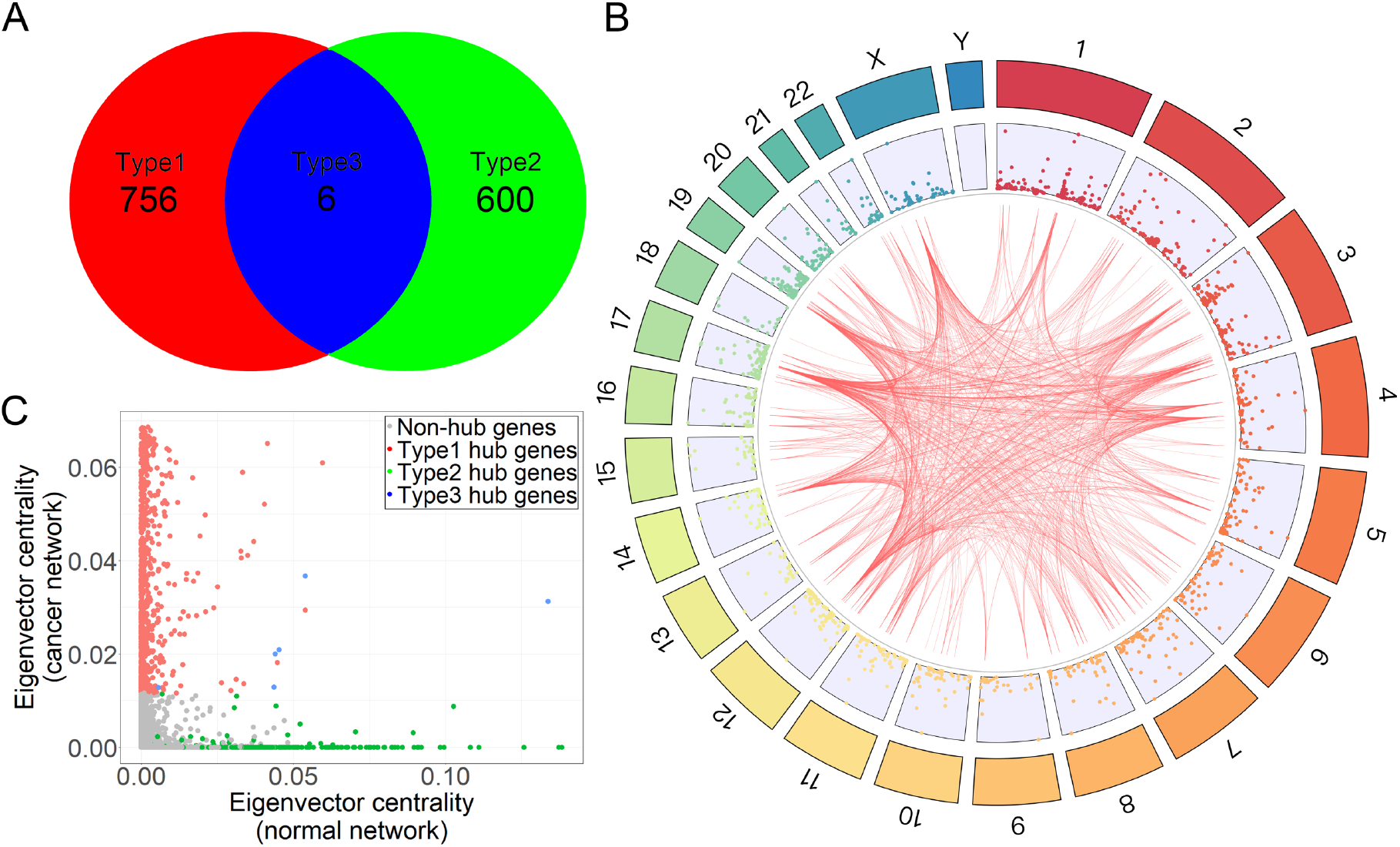
Cancer hub genes of the cancer gene relationship network. Type 1: hub genes enriched for only gained links; Type 2: hub genes enriched for only lost links; Type 3: hub genes enriched for both gained and lost links. (A) The number of three different types of cancer hub genes. (B) The distribution of cancer hub genes among chromosomes. The links inside the circle are the top 1000 links between cancer hub genes based on the NECARE output scores. (C) The centrality eigenvector of cancer hub genes. The x-axis is the centrality in the normal network, and the y-axis is the centrality in the cancer network.

Then, we classified cancer hub genes into three types: Type 1, hub genes enriched with gained links; Type 2, hub genes enriched with lost links; and Type 3, hub genes enriched with both gained and lost links. Overall, we identified 756 Type 1 hub genes, 600 Type 2 hub genes and 6 Type 3 hub genes (Fig. 3A). Fig. 3B shows their distribution among chromosomes and the top 1000 links with highest RI among all the hub genes. Even among hub genes, the top 1000 links were not distributed evenly, and some hub genes had more links than others. For example, CDK4 was engaged in 150 links among the top 1000 links and EGF was engaged in 109 links. In contrast, 39 hub genes engaged in only one link among the top 1000 links.

Type1 and Type2 hub genes were found enriched in very different pathways. Type 1 hub genes which tend to get new PPIs in cancer network were enriched in a lot of famous oncogenic signaling pathways (30), including: MAPK signaling pathway (P-value = 4.3×10-30), PI3K-Akt signaling pathway (P-value = 4.55×10-19), Ras signaling pathway (P-value = 1.56×10-14) and Wnt signaling pathway (P-value = 8.01×10-21) (Fig. S4A). Many famous cancer genes were Type1 hub genes including BRCA1, CDK1, CDK4, CDK14, CDKN1B, EGF, JUN, KRAS, MYC, YAP1. Meanwhile, Type 2 hub genes which tend to lose PPIs in cancer network were enriched in pathways for more general functions, such as Ribosome biogenesis in eukaryotes and Splicesome. One of the well-known Type 2 hub gene was TP53 (113 interactions lost, Table S5), which was correspond to the annotation from KEEG database (Fig. 1C). Besides, the most interesting result was that the type 2 hub genes were enriched in COVID19 pathway (Fig. S4B). This could be a kind of explanation of the previous finding that having cancer was an independent risk factor for in-hospital death from COVID-19 (31).

6 genes were Type 3 hub genes that had both gained- and lost-link perturbations (Fig. 3A, Table S6), including DYRK1A, LIMD1, POLR2B, S100A2, UBE2K and USH2A. Among them, DYRK1 is a dual kinase that can phosphorylate its own activation loop on tyrosine residue and phosphorylate its substrates on threonine and serine residues. It involves in different cancer pathways such as transcription, stress, DNA damage repair, apoptosis. And DYRK1A has the characteristic of duality in tumorigenesis. Some studies indicate its possible role as a tumor suppressor gene; however, others prove its pro-oncogenic activity (32). S100A2, which involves a number of cellular processes such as cell cycle progression and differentiation, was dysregulated in lung, gastric, esophageal, ovarian, bladder, breast, thyroid, melanoma and pancreatic cancer (33). And, LIMD1 was associated with head and neck squamous cell carcinoma, lung, gastric and breast cancer (34).

More interestingly, over 42% of genes that were found to be involved in cancer treatment were cancer hub genes in our study. Among them, 38% were Type 1 hub genes, 4% were Type 2 hub genes, and 4% were Type 3 hub genes. In addition, the distribution of the gained edges had no difference between clinically related genes and the background (all genes) (Kolmogorov–Smirnov P-value = 0.838, Fig. S4A). However, there was a significant difference in the distribution of the lost edges (Kolmogorov–Smirnov P-value < 8.5×10^-10^, Mean_All genes_ = 125 and Mean_Clinically related genes_ = 364) (Fig. S4B). Furthermore, those hub genes were significantly associated with the 10-year survival outcomes of 32 distinct types of cancer (Fig. 4). Overall, patients with high mutation scores had a poor prognosis and low survival rate (red lines in Fig. 4).

**Fig. 4:**
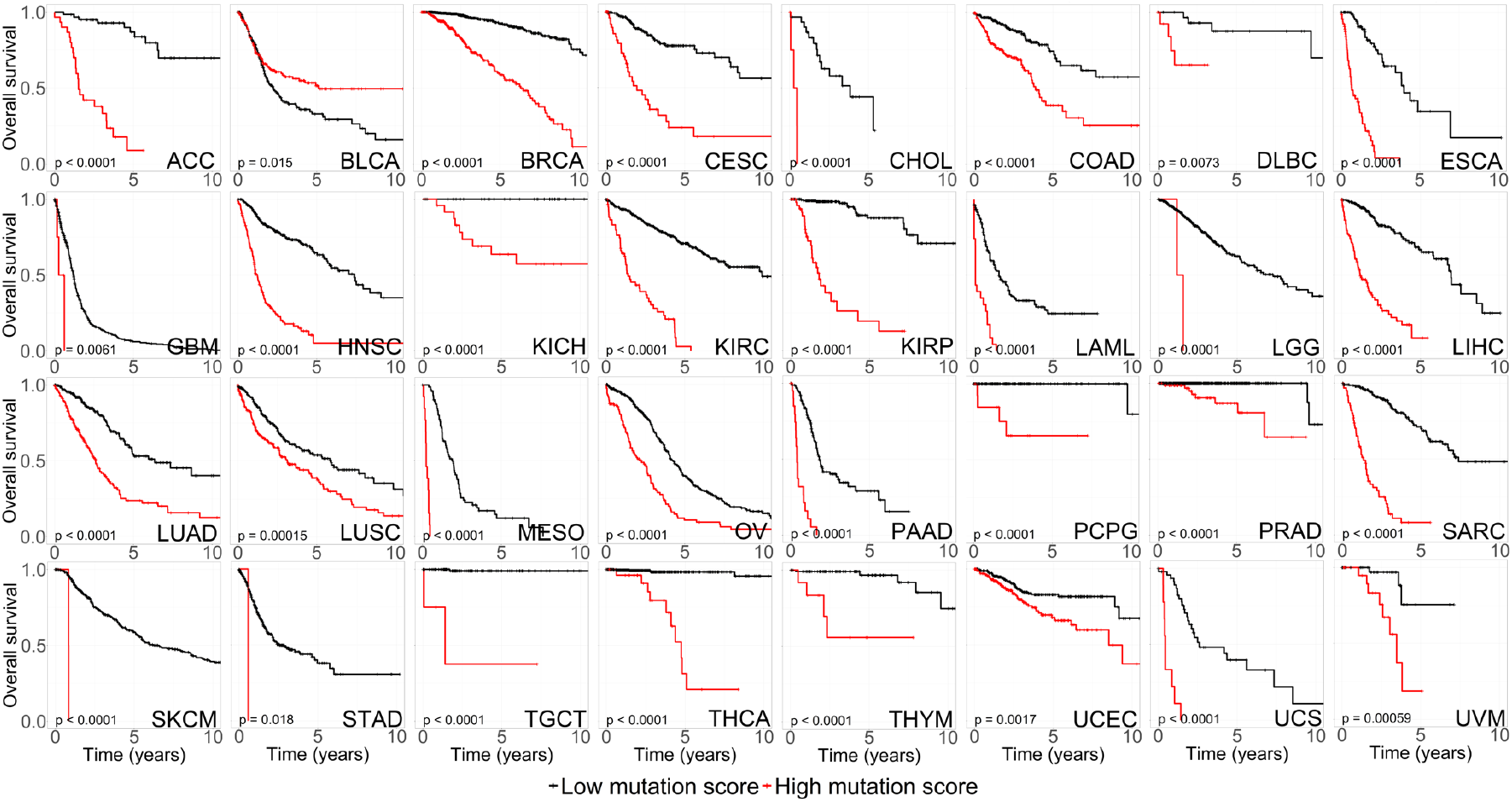
The prognostic landscape of hub genes. Kaplan–Meier plots for the patients from 32 different types of cancers from TCGA divided into high- and low-MS groups (Methods). The P-value was calculated by the log-rank test.

Subsequently, we analyzed the centrality of those hub genes (Fig. 3C). Three types of hub genes and the nonhub genes could be clearly separated by the centrality. First, this suggested that our statistical analysis, which was applied to identify hub genes, was reliable because we did not consider centrality during the identification of genes. Then, we found that Type 1 hub genes tended to have a high centrality in the cancer network but a low centrality in the normal network. However, Type 2 hub genes showed the reverse trend (a high centrality in the normal network but a low centrality in the cancer network). Type 3 hub genes were balanced between Type 1 and Type 2 hub genes. These nonhub genes had a low centrality in both normal and cancer networks. The centrality changes in Type 1 and Type 2 hub genes also reflected the perturbation of the cancer network.

### Experimental validation of NECARE predictions

The Ras signaling pathway is one of the most important pathways in cancer. Considering that pathways with too many genes (> 200) might be too generic, the Ras signaling pathway was filtered out and not discussed in our cross-talk analysis. However, there were clearly cross-talks between the Ras signaling pathway and other pathways. For example, there is cross-talk between the Wnt and Ras signaling pathways. Fig. 5A shows 10 highly reliable (RI > 90, Fig. 2A) bidirected gene interactions predicted by NECARE between WNT3 (from the Wnt signaling pathway) and SHC2 (from the Ras signaling pathway) with the following genes: RSPO4, CDK19, NR4A1, CDK8, AREG, LHX1, VGFR3, MAPK3, ZN619 and FGF9. WNT3 is a member of the Wnt family and may play a key role in cancer through activation of the Wnt-beta-catenin-TCF signaling pathway (35). SHC2 is located very upstream of the Ras signaling pathway and could be activated by many receptor tyrosine kinases (RTKs) in the Ras signaling pathway (6) (Fig. 5A).

**Fig. 5:**
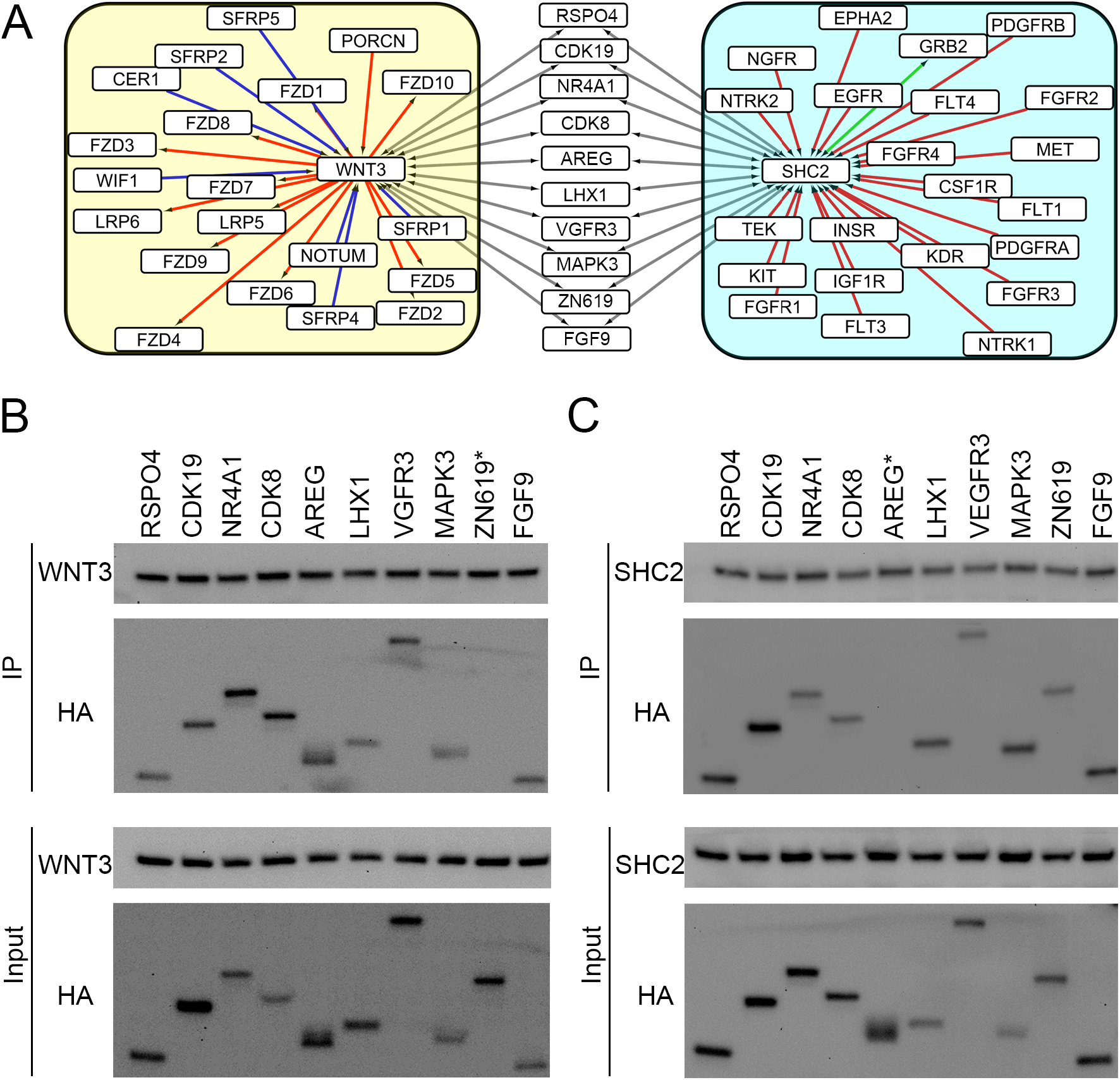
Experimental validation of the NECARE predictions. **Panel A** shows the genes that cross-talk with WNT3 and SHC2 in each pathway. Different colored edges represent different types of interactions. The red edge indicates activation; the blue edge indicates inhibition; the green edge is the KEGG annotated binding; the gray edge is NECARE predicted binding. The left yellow group shows the genes interacting with WNT3 in the Wnt signaling pathway. The right cyan group shows the genes in contact with SHC2 in the Ras signaling pathway. Those 10 genes in the middle with gray edges are NECARE predicted genes binding to WNT3 and SHC2 with a high RI (> 90). **Panels B** and **C** are co-IPs that validated the interactions of 10 predicted genes with WNT3 and SHC2 in LN229 cells. The interactions were determined by immunoblotting. The labelled “*” indicates a negative result of the co-IP validation experiment. **Panel B:** LN229 cells were co-transfected with the indicated HA-tagged constructs of 10 predicted genes and FLAG-tagged WNT3. **Panel C:** LN229 cells were co-transfected with the indicated HA-tagged constructs of 10 predicted genes and FLAG-tagged SHC2.

As bidirected positive predictions in NECARE were most likely to be the “binding” relationships, we applied coimmunoprecipitation (co-IP) to validate the predictions (SOM: Section S1, coimmunoprecipitation). We co-transfected the expression vectors of these 10 genes together with WNT3 and SHC2 in glioblastoma cell line LN229 (Fig. 5B, C). Co-IP was applied to confirm their binding interaction. 90% (18 of 20) of NECARE predictions were confirmed (Fig. 5B, C). Only two pairs of interactions, ZN619-WNT3 and AREG-SHC2, obtained negative validation results in co-IP (Fig. 5B, C).

## Discussion

Previous studies have already found several gene network perturbations in cancer. These results indicated that the gene network in cancer was significantly different from that in non-disease situations. In our study, we used R-GCN to establish a prediction method, NECARE, which is specific for cancer.

In the biological cell system, instead of isolation, genes act as a complex network. Genes may be regulated by others, control the expression of many other genes, or function together with other genes. Our model simulated this biological system by using a GCN. Information for each gene can be obtained from their neighbors in the network with weights for different relationship types and directions. Then, we compared our method with two different kinds of other algorithms: 1) sequence-based methods and 2) network-based methods. Our system outperformed all other algorithms in the task of predicting gene relationships in cancer. Sequence-based, state-of-the-art methods, such as PPI-Detect and PIPR (36), achieved good performance in PPI prediction of non-disease condition but failed in our cancer-specific task. Since genes were acting as a network complex, the disorder information would be broadcasted among the network. And the interaction between two genes may also be affected by their neighbors in the network. Therefore, sequence-based methods which only considered the input genes themselves may not be very specific for cancer gene relationship prediction. This is also the reason why we used network-based algorithm combined with knowledge-based features such as OPA2Vec. Our system with R-GCN can distinguish the relationship types and directions, while other network-based algorithms are not able to do so. Thus, our method is currently the best solution for cancer gene network prediction.

With the help of NECARE, we identified 1362 cancer hub genes that were enriched with network perturbations in cancer. As gene network perturbation was already found to be the main reason for cancer, these cancer hub genes should be the focus of the pathological mechanisms and treatment targets. Indeed, we found that a high mutation score of hub genes was significantly related to a poor prognosis of 32 different types of cancers. Over half of the cancer treatment-related genes in the database TARGET were hub genes in our study. Thus, these hub genes we identified have a high potential to be the drug design targets for cancer treatment and the other clinical research.

In addition, as mentioned before, we classified the hub genes into three types: Type 1 (gained links), Type 2 (lost links), and Type 3 (both gained and lost links). Unexpectedly, a lot of famous cancer genes were Type1 hub genes, and previous clinical studies also focused more on these hub genes. This phenomenon may be corresponding to the fact that cancer cells have their special characteristics, like limitless replicative potential, sustained angiogenesis and tissue invasion and metastasis. Gained links of genes in the network will for sure lead to the new functions of the whole cellular system, which can in some extent explain the behavioral characters of cancer cells. This can also explain why previous clinical studies also focused more on these hub genes. Targeting the newly established PPI in cancer cells may inhibit the new functions obtained by them, which can further block the uncontrolled proliferation, migration and invasiveness of cancer cells. Actually, there are also some famous cancer related genes, which not only got a lot of new interactions but also lost some links with other genes in cancer network. These results were corresponding to the previous studies that, instead of the simple destruction, cancer mutations lead the reconstruction of the gene network (7). So as a new perspective of cancer research which may lead to a better understanding of the pathological mechanism of cancer, we should also focus on how the cancer genes reprogramming the gene network with both the links they lost and the new interaction they got. Maybe this will provide a treatment strategy for those intractable cancers.

Overall, in our study, we established the first cancer-specific gene relationship prediction method. With the help of our new method, we analyzed gene network perturbations in cancer and identified cancer hub genes. Our method provides a powerful tool for biology researchers and clinicians to find possible interacting partners of their input genes in cancer. They can also choose to focus their research on the cancer hub genes identified by our method to develop new targets for cancer treatment.

## Conclusions

In conclusion, the gene relationship network in cancer was found to be different from that in non-disease situations in many previous studies. Here, we provide the novel cancer-specific network-based gene relationship prediction method: NECARE. Using NECARE, we revealed cancer gene network perturbations. We identified cancer hub genes that were enriched in network perturbations. Finally, we showed an example of NECARE application and validated the predictions by biological experiments.

## Supporting information

SOM

## ACKNOWLEDEGMENT

Financial support was obtained from the program of the China Scholarships Council (CSC201606230244, CSC201708080070). This work was also supported by a grant from the Outstanding Leaders Training Program of Pudong Health Bureau of Shanghai (PWR12018-07) and the Key Discipline Construction Project of Pudong Health Bureau of Shanghai (PWZxk2017-23, PWYgf2018-05). We also want to thank Prof. Jingbin Yan and Dr. Luksa Popovic for helping revise the manuscript.

## Author statement

**Jiajun Qiu**: Conceptualization and Methodology, **Kui Chen**: Validation, **Chunlong Zhong and Sihao Zhu**: Writing - Review & Editing, **Xiao Ma**: Writing - Original Draft and Project administration

