## Supplementary material for "Network-based protein-protein interaction prediction method maps perturbations of cancer interactome": SOM

### Supporting online material for:
### Network-based protein-protein interaction prediction method maps perturbations of cancer interactome

### Material

#### Section S1: Experimental validation of NECARE.

##### Cell culture and reagents

The commonly used glioblastoma cell line LN229 (ATCC, USA) was cultured with Dulbecco's Modified Eagle Medium (DMEM) full medium (DMEM, 10% FBS, 0.1 mg/mL streptomycin, 100 U/mL penicillin, and 0.025 mg/mL amphotericin B) and was maintained in 5% $CO_2$ atmosphere at 37°C.

##### Plasmid constructs and transfection

FLAG-WNT3 and FLAG-SHC2 were cloned into the pcDNA5 expression vector. Meanwhile, HA-RSPO4, HA-CDK19, HA-NR4A1, HA-CDK8, HA-AREG, HA-LHX1, HA-VGFR3, HA-MAPK3, HA-ZN619 and HA-FGF9 were subcloned into the pcDNA3.1 vector. LN229 cells were grown to be 70-80% confluent in 6-cm dishes before transfection. Then, they were transfected with Lipofectamine 3000 (Invitrogen) following the manufacturer's protocol. Cells were typically analyzed 24-36 h posttransfection.

##### Western blot analysis

The protein content of cell lysates was determined by western blot, and approximately 30 µg of whole lysates was used for each sample. In brief, whole lysates were separated by SDS polyacrylamide gel electrophoresis (SDS-PAGE) and transferred onto a polyvinylidene fluoride (PVDF) membrane. After blocking in 5% milk for 30 min, PVDF membranes were incubated overnight at 4°C with the corresponding primary antibodies: anti-FLAG (1:1000, Abcam), anti-HA (1:1000, Abcam) and anti-tubulin (1:2000, Abcam). The primary antibodies were detected with their corresponding horseradish peroxidase-conjugated secondary antibodies. The protein signals were detected using the enhanced chemiluminescence (ECL) substrate (Biocompare).

##### Coimmunoprecipitation (co-IP)

For co-IP, cells were harvested, and whole cell lysates were prepared with lysis buffer (50 mM Tris HCl, pH 7.4, 150 mM NaCl, 1 mM EDTA, and 1% Triton X-100). Then, whole cell lysates were sonicated and centrifuged for 10 min at 4°C at 11000 rpm. Flag-tagged proteins were purified anti-FLAG M2 agarose affinity gel (Sigma-Aldrich). After 2 h of incubation at 4°C, the FLAG immunoprecipitates were washed four times in lysis buffer and eluted with 1 × SDS sample buffer by boiling at 95°C for 10 min. Precipitated proteins were analyzed by western blotting.

**Table S1. Summary of general gene relationship data**

| Relationship Type | Count |
|---|---|
| dephosphorylation | 5 |
| ubiquitination | 38 |
| repression | 48 |
| phosphorylation | 90 |
| state change | 110 |
| expression | 3196 |
| post transcriptional modification | 9971 |
| inhibition | 14797 |
| compound | 25070 |
| activation | 41232 |
| catalysis | 46337 |
| reaction | 91341 |
| binding | 319615* |
| Total | 551850 |

*For the genes which bind itself, we only count it once

#### Fig. S1: Cross-validation procedure

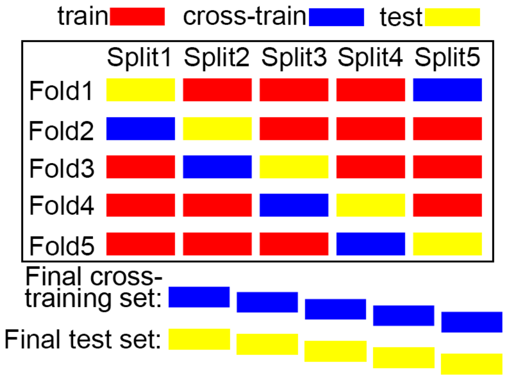

**Fig. S1: Cross-validation procedure.** For all machine learning developments, the original nonredundant data were split into five parts (Part 1-Part 5). Three parts were used for training, one for cross-training (optimization of hyperparameters, choice of feature), and one for testing. This was repeated five times (Fold 1-Fold 5, 5-fold cross-validation) so that each protein in the original data set had been used exactly once in the training set. Estimates for the standard error were compiled through bootstrap (Methods), not as the difference between the five folds.

### Results

**Fig. S2: Performance comparison among OPA2Vec alone, TCGA-based feature alone and the combination on the cross-training set.**

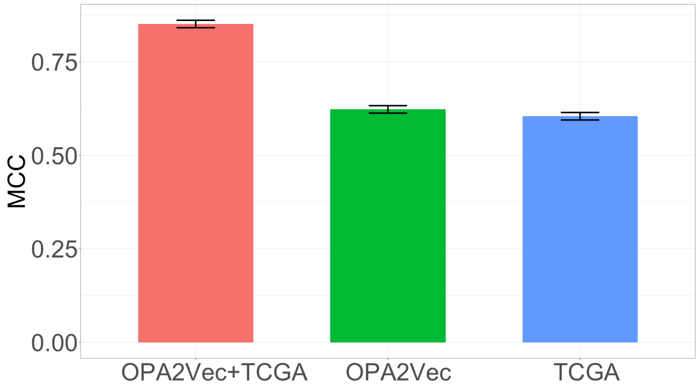

**Fig. S2: Performance comparison among OPA2Vec alone, TCGA-based feature alone and the combination on the cross-training set.** OPA2Vec means using the ontology-based feature OPA2Vec alone. TCGA means using only the TCGA-based expression and mutation profile. OPA2Vec+TCGA means using a combination of them.

#### Table S2 Cancer names in survival analysis from TCGA

| Study abbreviation | Study name |
|---|---|
| ACC | Adrenocortical carcinoma |
| BLCA | Bladder Urothelial Carcinoma |
| BRCA | Breast invasive carcinoma |
| CESC | Cervical squamous cell carcinoma and endocervical adenocarcinoma |
| CHOL | Cholangiocarcinoma |
| COAD | Colon adenocarcinoma |
| DLBC | Lymphoid Neoplasm Diffuse Large B-cell Lymphoma |
| ESCA | Esophageal carcinoma |
| GBM | Glioblastoma multiforme |
| HNSC | Head and Neck squamous cell carcinoma |
| KICH | Kidney Chromophobe |
| KIRC | Kidney renal clear cell carcinoma |
| KIRP | Kidney renal papillary cell carcinoma |
| LAML | Acute Myeloid Leukemia |
| LGG | Brain Lower Grade Glioma |
| LIHC | Liver hepatocellular carcinoma |
| LUAD | Lung adenocarcinoma |
| LUSC | Lung squamous cell carcinoma |
| MESO | Mesothelioma |
| OV | Ovarian serous cystadenocarcinoma |
| PAAD | Pancreatic adenocarcinoma |
| PCPG | Pheochromocytoma and Paraganglioma |
| PRAD | Prostate adenocarcinoma |
| SARC | Sarcoma |
| SKCM | Skin Cutaneous Melanoma |
| STAD | Stomach adenocarcinoma |
| TGCT | Testicular Germ Cell Tumors |
| THCA | Thyroid carcinoma |
| THYM | Thymoma |
| UCEC | Uterine Corpus Endometrial Carcinoma |
| UCS | Uterine Carcinosarcoma |
| UVM | Uveal Melanoma |

**Table S3 Optimized hyperparameter of NECARE in cross-training set**

| Name of parameters | Values |
|---|---|
| Learning rate[1] | 0.01 |
| Number of hidden layers | 2 |
| Number of hidden nodes | 200 |
| Number of bases[2] | 10 |
| Dropout | 0.2 |

[1]The learning rate for first 30 epoch was set as 0.1, and it was slow down to 0.01 after 30 epoch.

[2] Basis decomposition was applied to reduce model parameter size and prevent overfitting in RGCN(1). Number of bases would be a common divisor for the number of hidden nodes and the length of features. To make the optimization easier, we shrank the length of final features to be 200 through PCA.

**Table S4 Summary of the comparison based on test set**

| Method | AUC | Precision (%) | Recall (Sensitivity,%) | Specificity (%) | F1 (%) | MCC |
|---|---|---|---|---|---|---|
| NECARE | 0.97±0.02 | 90±2 | 92±2 | 92±2 | 90±2 | 0.84±0.03 |
| AA(2) | 0.71±0.02 | 51±1 | 94±2 | 35±1 | 66±1 | 0.33±0.03 |
| CAA(3) | 0.67±0.02 | 54±2 | 77±2 | 54±2 | 64±2 | 0.31±0.03 |
| CAR(3) | 0.67±0.02 | 56±2 | 75±2 | 58±2 | 64±2 | 0.33±0.03 |
| CN(3) | 0.72±0.02 | 51±1 | 93±2 | 36±1 | 66±1 | 0.33±0.03 |
| CPA(3) | 0.52±0.01 | 42±1 | 99±1 | 2±1 | 59±1 | 0.03±0.03 |
| CRA(3) | 0.67±0.02 | 55±2 | 78±2 | 54±2 | 65±2 | 0.33±0.03 |
| Dice index(4) | 0.68±0.02 | 51±1 | 94±2 | 36±1 | 66±2 | 0.34±0.03 |
| DLPPI(5) | 0.50 | 42±1 | 100 | 0 | 59±1 | 0.00 |
| DPPI(6) | 0.60±0.02 | 50±2 | 68±2 | 47±2 | 58±2 | 0.15±0.04 |
| Jaccard index(7) | 0.67±0.02 | 51±1 | 94±2 | 36±1 | 66±1 | 0.34±0.03 |
| L3(8) | 0.63±0.02 | 44±1 | 99±1 | 9±1 | 61±1 | 0.19±0.03 |
| MCE(9) | 0.48±0.02 | 40±1 | 88±1 | 5±1 | 55±1 | -0.12±0.03 |
| PIPR(10) | 0.94±0.06 | 90±5 | 84±5 | 83±8 | 86±5 | 0.65±0.10 |
| PPI-Detect(11) | 0.56±0.02 | 45±2 | 66±2 | 42±2 | 54±2 | 0.09±0.03 |
| Preferential Attachment(12) | 0.514±0.005 | 42±1 | 100 | 0 | 59±1 | 0.00±0.02 |
| RA(13) | 0.69±0.02 | 52±1 | 89±1 | 40±2 | 65±1 | 0.32±0.03 |

*Note*: ± gives the error estimated by bootstrapping (Methods). Percentages for precision, recall, specificity, and F1 , MCC score (Eqn. 3 and Eqn. 4)

#### Fig. S3: Distribution of gained or lost edges.

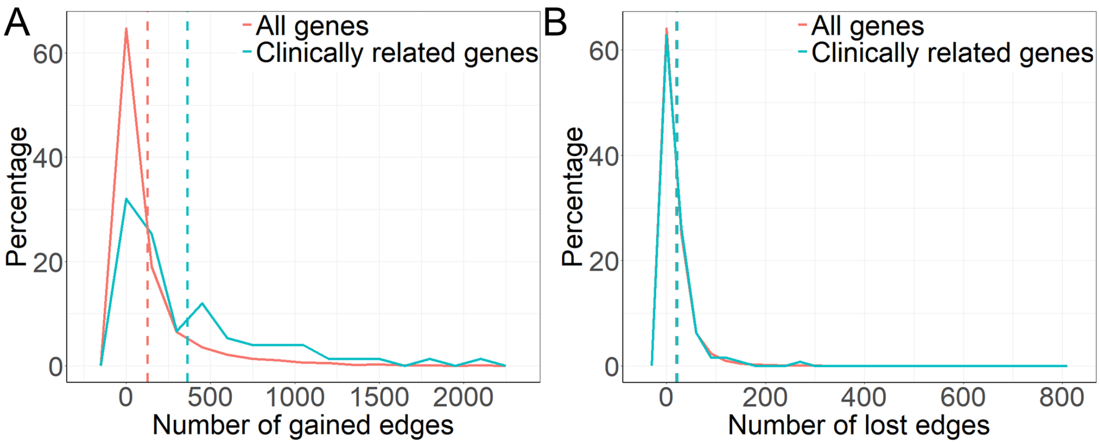

**Fig. S3: Distribution of gained or lost edges.** (A) The distribution of gained edges. The dashed lines represent the mean. $\text{Mean}_{\text{All genes}}$ = 125 and $\text{Mean}_{\text{Clinically related genes}}$ = 364. (B) The distribution of lost edges. The dashed lines represent the mean. $\text{Mean}_{\text{All genes}}$ = 20 and $\text{Mean}_{\text{Clinically related genes}}$ = 21.

#### Table S5 Cancer hub genes

(See independent file: Table S6.tsv)

#### Fig. S4: KEGG enrichment analysis for cancer hub genes.

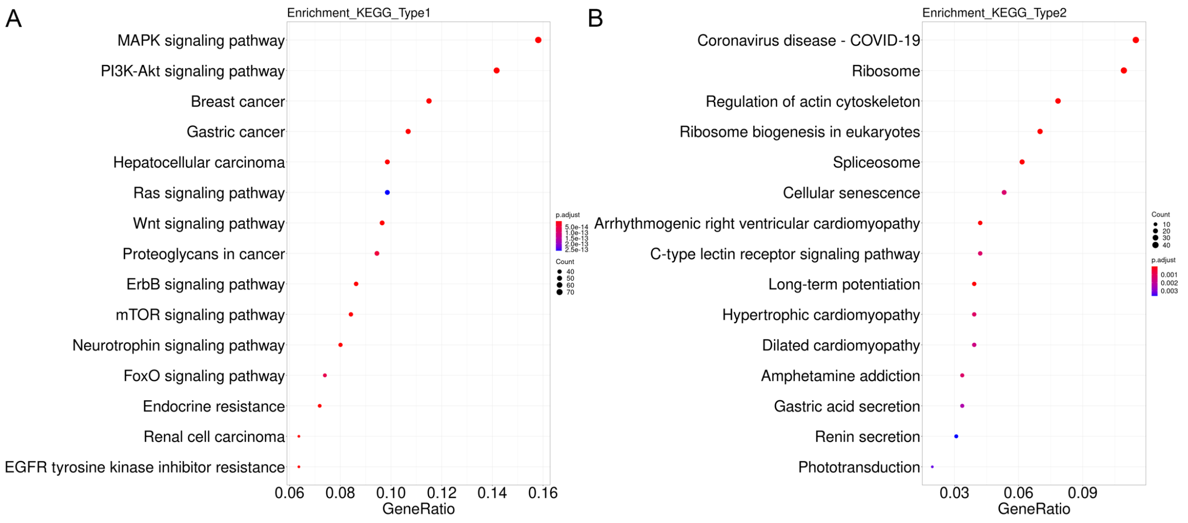

**Fig. S4: KEGG enrichment analysis for cancer hub genes.** The x-axis is the gene ratio, which represents the percentage of all genes annotated to a pathway. Dot size is the number of genes annotated to a pathway. The color of each dot corresponds to the P-value of KEGG enrichment analysis. (A) KEGG enrichment analysis for Type1 hub genes. (B) KEGG enrichment analysis for Type2 hub genes.
